# IKZF3 deficiency potentiates chimeric antigen receptor T cells targeting solid tumors

**DOI:** 10.1101/2021.06.18.449074

**Authors:** Yan Zou, Tian Chi

## Abstract

Chimeric antigen receptor (CAR) T cell therapy has been successful in treating hematological malignancy, but solid tumors remain refractory. Here, we demonstrated that knocking out transcription factor IKZF3 in HER2-specific CAR T cells targeting breast cancer cells did not affect proliferation or differentiation of the CAR T cells in the absence of tumors, but markedly enhanced killing of the cancer cells *in vitro* and in a xenograft model. Furthermore, *IKZF3* KO had similar effects on the CD133-specific CAR T cells targeting glioblastoma cells. AlphaLISA and RNA-seq analyses indicate that *IKZF3* KO increased the expression of genes involved in cytokine signaling, chemotaxis and cytotoxicity. Our results suggest a general strategy for enhancing CAR T efficacy on solid tumors.

## Introduction

Chimeric Antigen Receptor (CAR) T cells are human primary T cells genetically modified to express a CAR, the latter comprising the extracellular single-chain variable fragment (scFv) targeting surface-exposed tumor-associated antigens (TAA) in non-MHC restricted manner, a transmembrane domain, and one or more intracellular signaling domains that activate T cells upon antigen engagement^1^. CAR T cells are effective in eradicating some hematological malignancies^2-5^. However, despite much effort, solid tumors have proved largely unresponsive due to inefficient infiltration of CAR T cells, presence of immunosuppressive tumor microenvironment and exhaustion of T cells^1, 6, 7^. Various strategies have been attempted to potentiate CAR-T cells targeting solid tumors, including the expression of chemokine receptors^8-10^, disruption of the inhibitory adenosine and TGF-β signaling via deletion of adenosine and TGF-β receptor *A2AR* and *TGFBR2*, respectively^11, 12^, administration of IL-7, IL-15 or IL-21^13-15^, use of immune checkpoint inhibitors^16-18^, or deletion of PD-1 gene in CAR T cells^19-21^, but their benefits are limited. For example, the use of immune checkpoint inhibitors in conjunction with CAR T therapy may increase the risk of autoimmunity, while PD-1 knockout may lead to the failure of CD8+ T cells, as PD-1 expression can protect CD8+ T cells from excessive proliferation and terminal differentiation^22^. It is thus highly desirable to devise additional methods to harness the power of CAR T cells for solid tumor treatment.

Lenalidomide (LENA) is an immunomodulatory drug with pleotropic effects on diverse immune cells including natural killer cells, monocytes, B cells and T cells^23-25^. In T cells, LENA can increase IL-2 induction upon TCR stimulation^26^, shift T helper responses from Th2 to Th1^27^, and inhibit Treg function^28^. LENA acts mainly by causing the ubiquitin-mediated degradation of the Ikaros family zinc finger transcription factor IKZF1 and IKZF3^29, 30^. In T cells, siRNA-mediated knockdown of either IKZF1 or IKZF3 is sufficient to enhance the expression of IL-2 and multiple other cytokines, reminiscent of the effects of LENA treatment^29^. We have recently demonstrated that *in vitro*, LENA can enhance cytokine expression and killing of solid tumors by CAR T cells, suggesting that LENA might be used for boosting the efficacy of CAR T cells targeting solid tumors^31^. However, LENA has been FDA-approved only for treating several hematological cancers (namely multiple myeloma, myelodysplastic syndromes and mantle cell lymphoma), but not for any solid tumors. One potential strategy to overcome this barrier would be to create CAR T cells deficient in IKZF1 and/or IKZF3, which may recapitulate the beneficial effects of LENA on T cells but can avoid the unintended pleiotropic effects on other cell types that would result from systemic administration of LENA. In this study, we have opted to target IKZF3 instead of IKZF1 in CAR T cells, because IKZF1 has the characteristics of a tumor suppressor gene: IKZF1 can arrest the uncontrolled growth and proliferation when introduced into an established mouse Ikaros null T-leukemia cell line^32-34^, and IKZF1 promotes the development of myeloproliferative neoplasms^35^. In contrast, such effects have not been described for IKZF3, suggesting it is safer to knock out IKZF3 in T cells as compared with IKZF1. Our results demonstrate that *IKZF3* KO markedly enhanced the ability of CAR T cells to kill solid tumor cells *in vitro* and in a xenograft model, which was associated with increased expression of genes mediating cytokine signaling, cell trafficking and cytotoxicity. These data suggest a novel strategy for boosting the efficacy of CAR T therapy for solid tumors.

## Results and Discussion

### Generation of IKZF3-deficient HER2-specific CAR T cells

The CAR T cells we sought to optimize expressed HER2-targeting CAR comprising a Myc-tagged scFv derived from the Herceptin monoclonal antibody, the CD8 transmembrane domain upstream of the tandem signaling domains from CD28, 4-1BB and CD3-ζ^36^, which is a commonly used form of CAR^37^. To generate IKZF3-deficient HER2-CAR T cells (termed HER2K3-CAR T), we nucleofected PBMCs from two healthy donors with expression vectors for CAR, piggyBac transposase, Cas9 and the dual gRNAs targeting IKZF3 exon3; the dual gRNA format was used due to higher editing efficiency^38^. Control CAR T cells (termed HER2-CAR T) were generated in parallel by omitting Cas9 and gRNA expression vectors. Nucleofected cells were then cultured with irradiated allogenic PBMCs, anti-CD3 antibody, IL-2 and puromycin to expand and select CAR-T cells. The cells at the end of the expansion were then characterized.

We found that over 89% and 79% of the Cas9/gRNA transfected CAR T cells from Donor 0529 and Donor 0710, respectively, harbored out-of-frame mutations at IKZF3 exon 3 (Figure 1A) concomitant with marked reduction in IKZF3 protein expression (Figure 1B), thus demonstrating highly efficient editing. Proliferation of these cells was somewhat impaired by IKZF3 disruption (Figure 1C) but importantly, CAR expression was unaffected (Figure 1D-E), and neither was the phenotype related to cell differentiation altered as defined by CD62L and CD45RO expression (Figure 1F-G; note that donor-specific differences in the relative abundance of central vs effector memory cells). Thus, IKZF3 was dispensable for CAR expression or cell differentiation during the expansion and selection of the CAR T cells. However, IKZF3 deficiency induced CD69 expression in CAR T cells cocultured with allogeneic PBMCs (Figure 1H-I), indicating IKZF3 involved in TCR stimulation signals downstream of allogeneic response.

**Figure 1.**
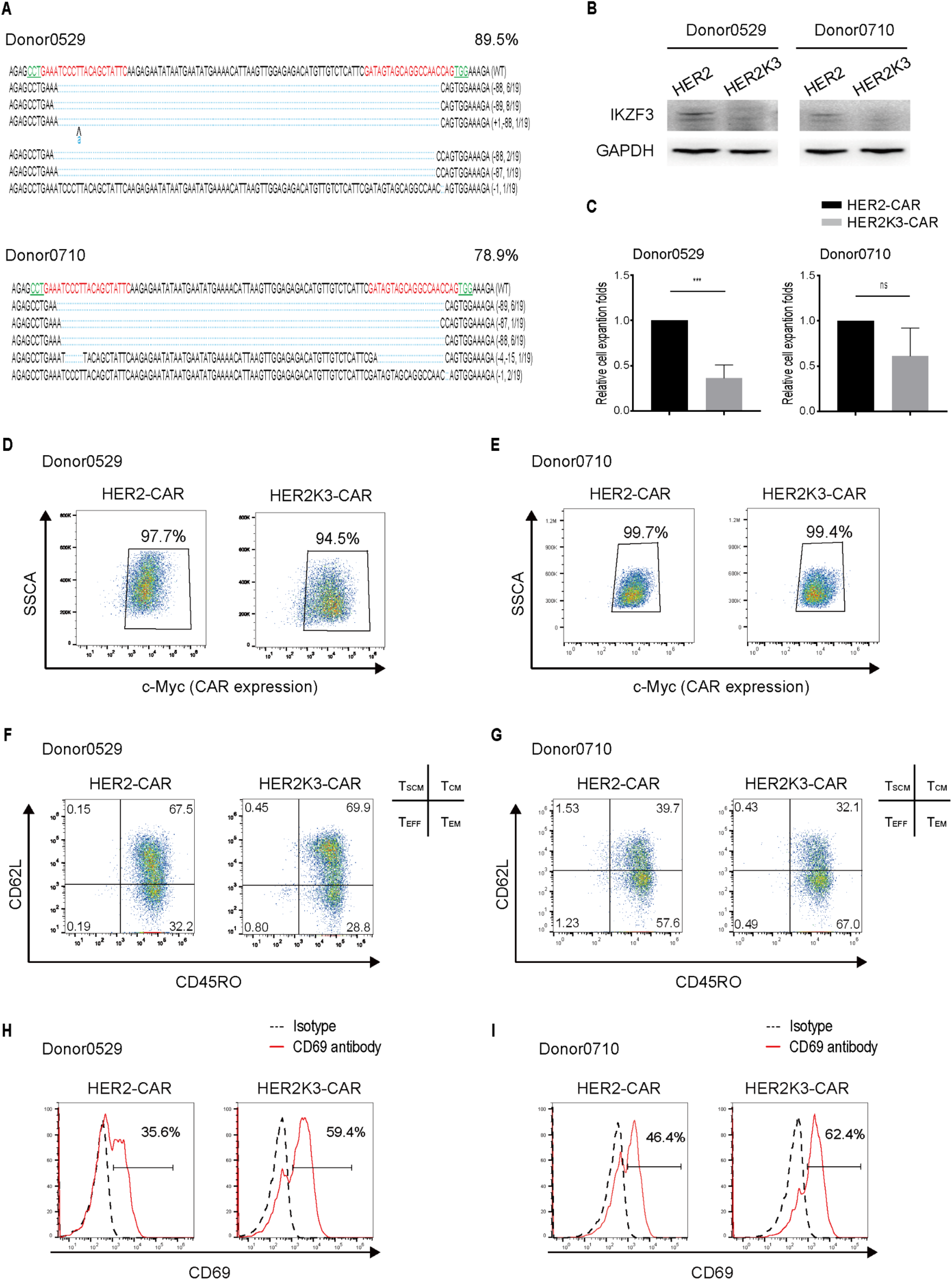
Basic features of *IKZF3* KO CAR T cells targeting HER2. (A) Successfully edited *IKZF3* alleles in the CAR T populations. For CAR T cells derived from each donor, the targeted region was PCR-amplified and TA-cloned before 19 bacterial colonies were picked for Sanger sequencing. The values in the brackets indicate the numbers of deleted/inserted base pairs and the frequencies of the occurrence of the alleles among the 19 samples. PAM sites for the dual gRNAs are highlighted. (B) Western blot showing reduced IKZF3 expression in the crude edited population. (C) Relative CAR T cell numbers at the end of the expansion. Values are mean +/- SD (n= 3 technical repeats, \**p*<0.05, \*\**p*<0.01, \*\*\**p*<0.001). (D-G) FACS analysis showing little effect of *IKZF3* KO on CAR expression (based on anti-Myc staining; D-E) and CAR T cell subset composition (based on CD45RO and CD62L expression; F-G). (H-I) Increased CD69 expression frequency in KO CAR T cells.

### IKZF3 deficiency potentiates killing of breast cancer by HER2-specific CAR T cells

We next investigated the cytolytic efficacy of the HER2K3-CAR T cells using three assays.

First, the CAR T cells were incubated with MDA-MB-453 cells (originating from a human mammary gland cancer) expressing firefly luciferase (termed “453-ffluc” hereafter) for 48 h, and the loss of luciferase activity within viable cells was taken as a measure of cell lysis. HER2K3-CAR T cells showed enhanced killing compared with HER2-CAR T cells (Figure 2A), which was correlated with augmented production of IFN-γ, IL-2, TNF-α and GM-CSF (Figure 2B). We also determined the proliferation of the CAR T cells by culturing CFSE-labeled CAR T cells with irradiated MDA-MB-453 cells. After seven days of culture, the HER2K3-CAR T population harbored more divided cells than HER2-CAR T (78.5% vs. 62.7%) and had undergone more rounds of division (1.4 vs 1; Figure 2C), which correlated with higher frequency of CD69 expression (Figure 1H-I), indicating that IKZF3 deficiency facilitated tumor-stimulated CAR T cell activation and proliferation.

**Figure 2.**
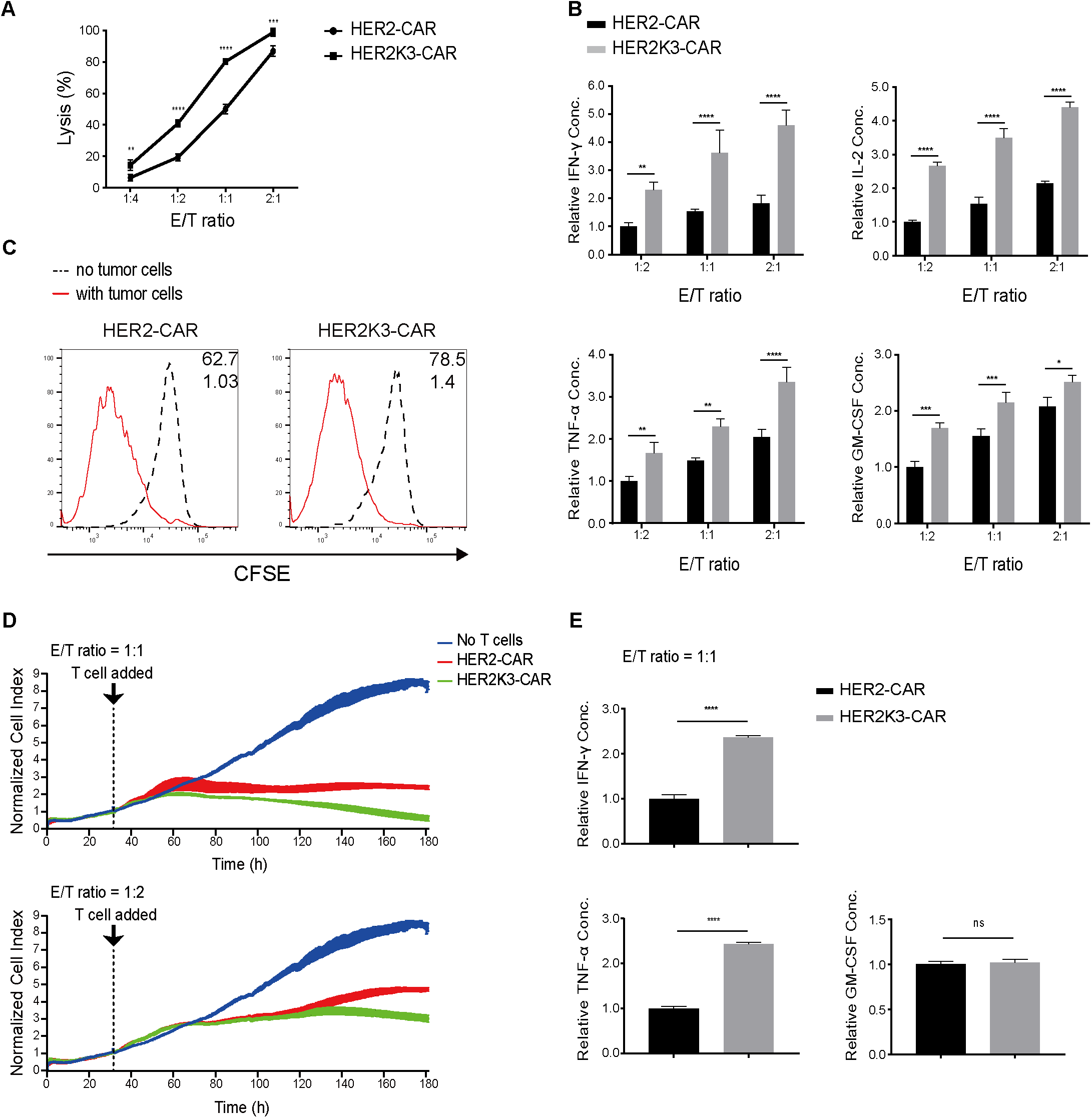
*IKZF3* KO enhances HER2-CAR T activity *in vitro*. (A) Cytolysis of the target cells (luciferase-expressing MDA-MB-453) after 48 h co-incubation with CAR T cells at indicated effector to target (E/T) ratios. (B) Relative cytokine concentration in the culture media after 24 h co-incubation. (C) Proliferation of CFSE-labeled CAR T cells after 7-day culture in the presence (red) or absence (black) of lethally irradiated MDA-MB-453 cells. The values inside the plots are the “Percent Divided” (top) and the “Division Index” (bottom; see Materials and Methods). (D) Real time monitoring of cytolysis using impedance-based assay for 180 h. MDA-MB-453 cells, attached to the bottom of electrode plate, were cultured with CAR T cells at 1:1 (top) or 1:2 (bottom) effector to target ratio, and cell index (reflecting changes in impedance caused by the loss of tumor cells) measured every 15 min. (E) Cytokine concentration in the media at the end of 180 h incubation. Values are mean+/- S.D (n=3 technical repeats). Groups were compared through two-way ANOVA or two-tailed unpaired t-test. \**p*<0.05, \*\**p*<0.01, \*\*\**p*<0.001, \*\*\*\**p*<0.0001. IL-2 was undetectable (not shown).

Second, impedance-based killing assays were carried out to assess the cytolytic activity over a longer period of time. Specifically, MDA-MB-453 cells were attached to the electrode plate, and CAR T cells then added to lyse and detach the tumor cells, which was read out as changes in the electrical impedance. We measured the impedance every 15 min over a 150 h period, and also determined cytokine concentration in the media at the end of the experiment. HER2K3-CAR T cells were more effective at tumor killing at various effector to target ratios over the entire course of the experiment (Figure 2D), correlated with higher IFN-γ and TNF-α concentration at the end of the experiment, although by this time, GM-CSF concentrations had become comparable in both groups of T cells while IL-2 had become undetectable (Figure 2E and data not shown). The same trend was observed for the CAR T cells derived from the second donor (Figure S1).

Finally, we assayed tumor killing *in vivo*. Immunodeficient NPSG mice were subcutaneously inoculated with 1.5 ⨯ 10^6^ 453-ffluc cells. Six days later, 5 ⨯ 10^6^ CAR T cells were injected i.v. and tumor growth monitored using bioluminescence imaging at various times thereafter. HER2K3-CAR T cells proved more efficient at tumor suppression, with bioluminescence eliminated in the 3 of 5 mice (as opposed to 1 out of 5 mice for HER2-CAR T; Figure 3A, 3B), leading to enhanced mice survival (Figure 3C).

**Figure 3.**
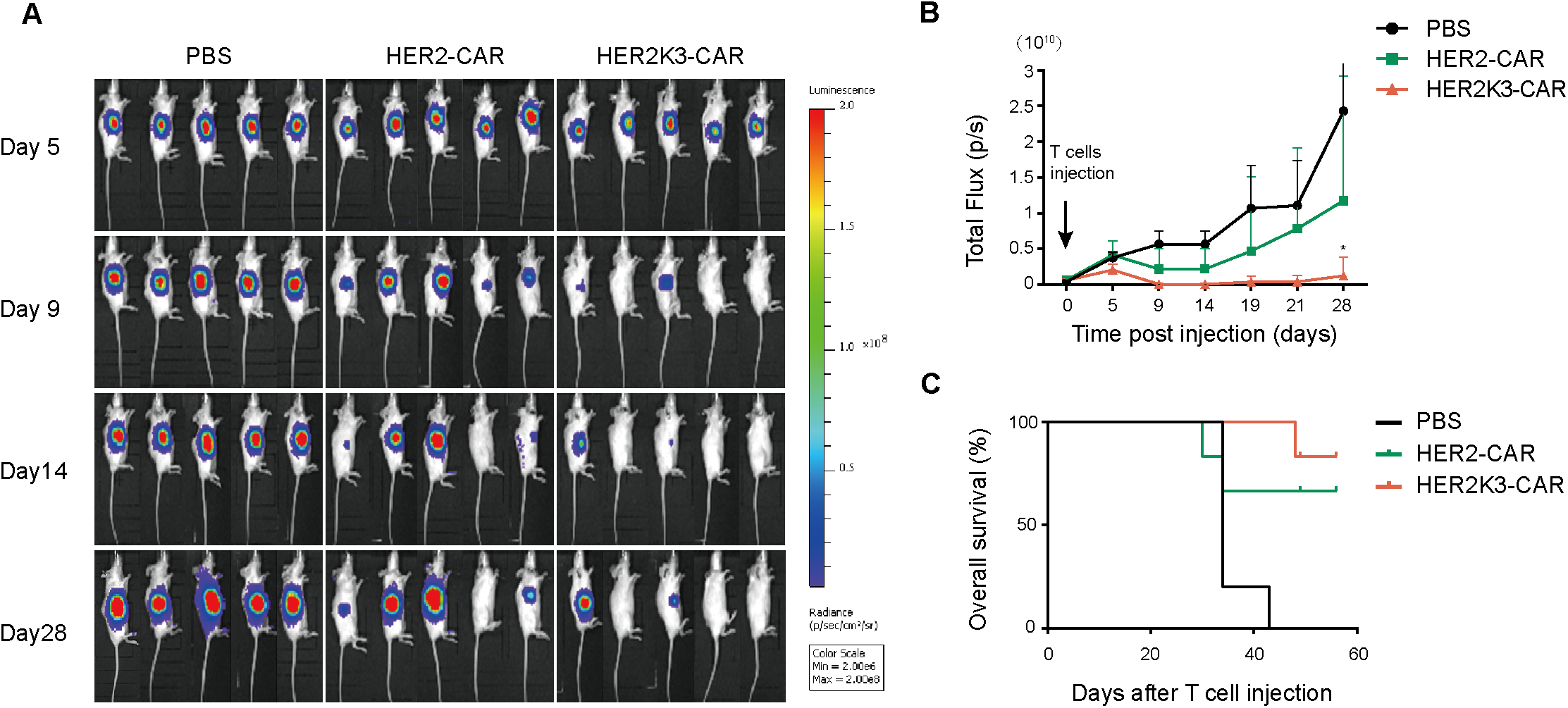
*IKZF3* KO enhances HER2-CAR T activity *in vivo*. Firefly luciferase-expressing MAD-MB-453 cells (1.5 × 10^6^) were inoculated subcutaneously in the dorsal flank of right foreleg of the NPSG mice (5 mice/group) before intravenous injection of CAR T cells (5 × 10^6^) 6 days later. Bioluminescence was imaged at various times thereafter (A), the luminescence intensity was quantified (B), and mice survival recorded (C). Groups were compared through two-way ANOVA. Error bars, s.d.; \**p*<0.05.

We conclude that HER2K3-CAR T cells were more potent than HER2-CAR T cells at killing MDA-MB-453 tumor cells, which might result from enhanced activation, proliferation and cytokine production following tumor stimulation.

### IKZF3 deficiency upregulates genes important for T cell function

To further understand how IKZF3 deficiency augmented CAR T cell function, we used RNA-seq to compare gene expression patterns in HER2-CAR T vs. HER2K3-CAR T cells, finding *IKZF3* KO upregulated 106 genes and downregulated 92 genes (excluding genes encoding TCR variable regions; Fig. 4A), which contained 97 and 86 protein-coding genes, respectively (Supplementary Table).

**Figure 4.**
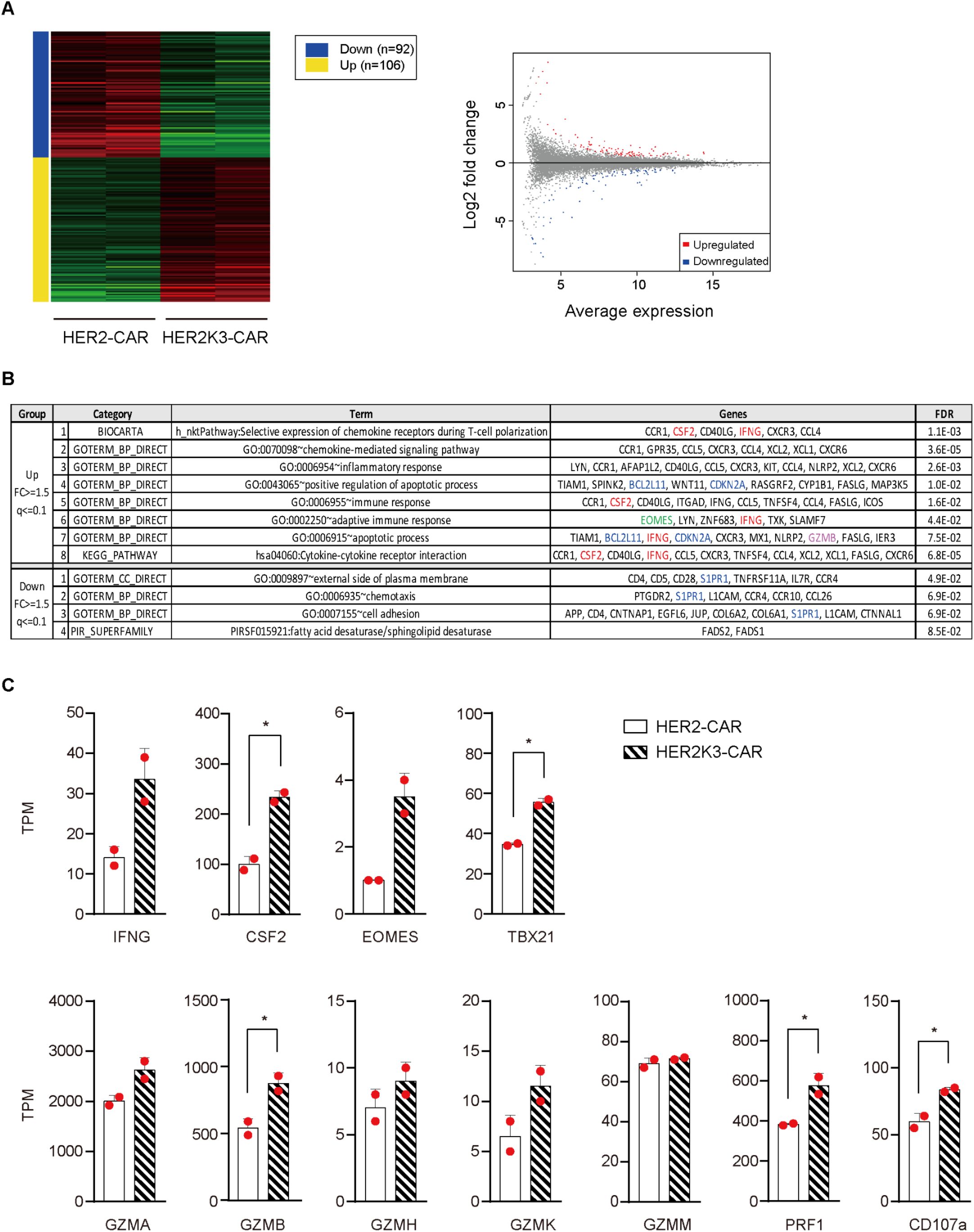
Transcriptome changes caused by *IKZF3* KO. (A) Differentially expressed genes (DEGs) revealed by RNA-seq, shown in heatmap (left) and MA-plot (right). DEGs are defined as the genes whose expression are changed by at least 1.5x (q<=0.1). RNA-seq was performed twice on different dates, using the CAR T cells derived from Donor 0710 (biological replicates). (B) Enriched pathways in the upregulated (top) and downregulated (bottom) DEGs, as analyzed by DAVID. GO, Gene Ontology; KEGG, Kyoto Encyclopedia of the Genes and Genome; FDR, False Discovery Rate from Benjamini and Hochberg. (C) Expression levels (in TPM) of selected genes from RNA-seq data. The values are mean+/- SD (n=2 biological replicates).

Multiple biologically interesting pathways were detected among the 97 upregulated genes, which jointly suggested that *IKZF3* KO enhanced cytokine signaling, chemotaxis, adhesion and immune responses, consistent with the potentiation in the killing ability (Fig. 4B). IFNG and CSF2/GM-CSF were repetitively enriched in multiple pathways (Categories 1, 5-8, red), suggesting their functional importance. The extents of the upregulation of IFNG and CSF2/GM-CSF in the KO cells, as revealed by manual examination of the RNA-seq data, were moderate (2-3 folds, Fig. 4C, top left), echoing the results of the AlphaLISA assay (Fig. 2B). In addition, the transcription factor Eomes was also upregulated (Fig. 4B, Category 6), as was TBX21 (Fig. 4C, top right). Both factors are important regulators of CD8 T cell function^39^, with Eomes upregulated in effector CD8 cells and contributing to IFN-γ production and cytotoxicity of CD8 T cells^40, 41^, while TBX21 is upregulated and plays important roles in memory CD8 T cells. It is also noteworthy that genes capable of apoptosis regulation were also upregulated (Categories 4, 7), which included GZMB important for T cell cytotoxicity (Fig. 4B, pink gene in Category 7). Remarkably, manual examination of the RNA-seq data indicates that the remaining members of GZM family, namely GZMA, GZMH, GZMK and GZMM, all tended to be overexpressed in the KO cells except GZMM (Fig. 4C, bottom). Furthermore, Perforin and the degranulation marker CD107a were also upregulated (Fig. 4C, bottom). Thus, enhanced killing might also result from overexpression and degranulation of granzymes and perforin in CD8 T cells. Paradoxically, the apoptosis-related genes overexpressed in the KO cells included the proapoptotic genes BCL2L11 and CDKN2A, with unclear biological relevance (Categories 4 and 7, blue).

In contrast to the upregulated genes, for the 86 downregulated genes, all pathways detected had relatively large FDR (>0.049), and only 4 pathways had FDR<0.1 (Fig. 4B, bottom). The relevance of these 4 enriched pathways to CAR T function was unclear. Of note, S1PR1, a pro-survival gene, was present in 3 out of the 4 pathways (Fig. 4B, bottom, blue). S1PR1 downregulation reinforced the notion that the KO cells might be apoptotic, as first suggested by BCL2L11 and CDKN2A upregulation.

We conclude that IKZF3 deficiency in HER2-CAR T cells enhanced the expression of genes promoting cytokine signaling, chemotaxis and cytotoxicity, which might underlie the potentiation of CAR T therapeutic effects. However, IKZF3 deficiency might also cause a “side-effect”, namely an increase in CAR T apoptosis. Countering this putative effect could perhaps further improve the function of the KO CAR T cells.

### IKZF3 deficiency potentiates CD133-specific CAR T cells

Finally, we determined whether *IKZF3* KO could potentiate CAR T cells targeting a different antigen such as CD133, a biomarker for tumor-initiating cells in multiple human cancers, especially glioblastoma^42^, where a small population of CD133 positive glioblastoma tumor-initiating cells manage to survive radiotherapy and chemotherapy, conferring resistance to the therapies^43, 44^.

CD133-specific CAR (133-CAR) T cells and their IKZF3-deficient (133K3-CAR) counterpart were generated from two donors as in the case of the HER2-specific CAR T cells. In the 133K3-CAR T cells from both donors, the *IKZF3* gene was efficiently mutated (Figure 5A) and the protein eliminated (Figure 5B). *IKZF3*-deficiency did not affect cell expansion (Figure 5C) or CAR expression (Figure 5D, 5E), but for unknown reasons, enriched or partially depleted the CD62L^+^ CD45RO^+^ central memory T cell subset depending on the T cell donors (Figure 5F-G). CD69 expression was increased in *IKZF3* KO (Figure 5H-I) as indicated in HER2-CAR T cells (Figure 1H-I). Importantly, for both donors, *IKZF3*-deficiency potentiated killing of the glioblastoma U251 cells overexpressing CD133 (U251-OE-ffluc) in short-term cultures (Figure 6A; Figure S2A), concomitant with increased cytokine production (Figure 6B; Figure S2B), cell proliferation (Figure 6C) and CD69 induction (Figure 5H-I; Figure S2C). In long-term (160 h) cultures, *IKZF3*-deficiency similarly potentiated killing (Figure 6D; Figure S2D), which was associated with increased production of IFN-γ and TNF-α (but not IL-2 or GM-CSF; Figure 6E). Thus, the effects of IKZF3 elimination on CD133-specific CAR T cells were similar to that on HER2-CAR T cells.

**Figure 5.**
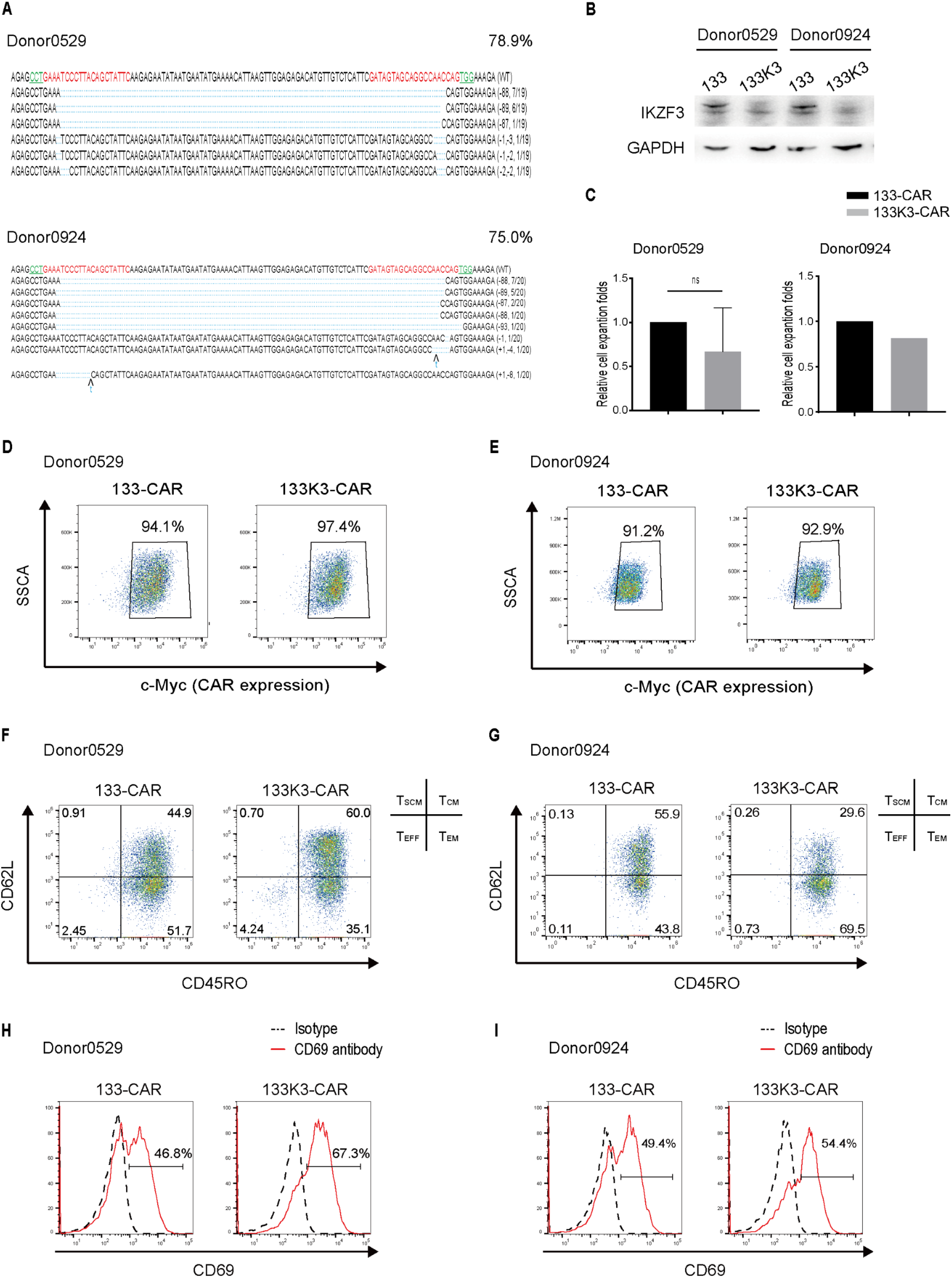
Basic features of IKZF3-deficient CD133-specific CAR T cells. (A-I) Same as Fig. 1A-I, except that the CAR T cells targeting CD133 were analyzed

**Figure 6.**
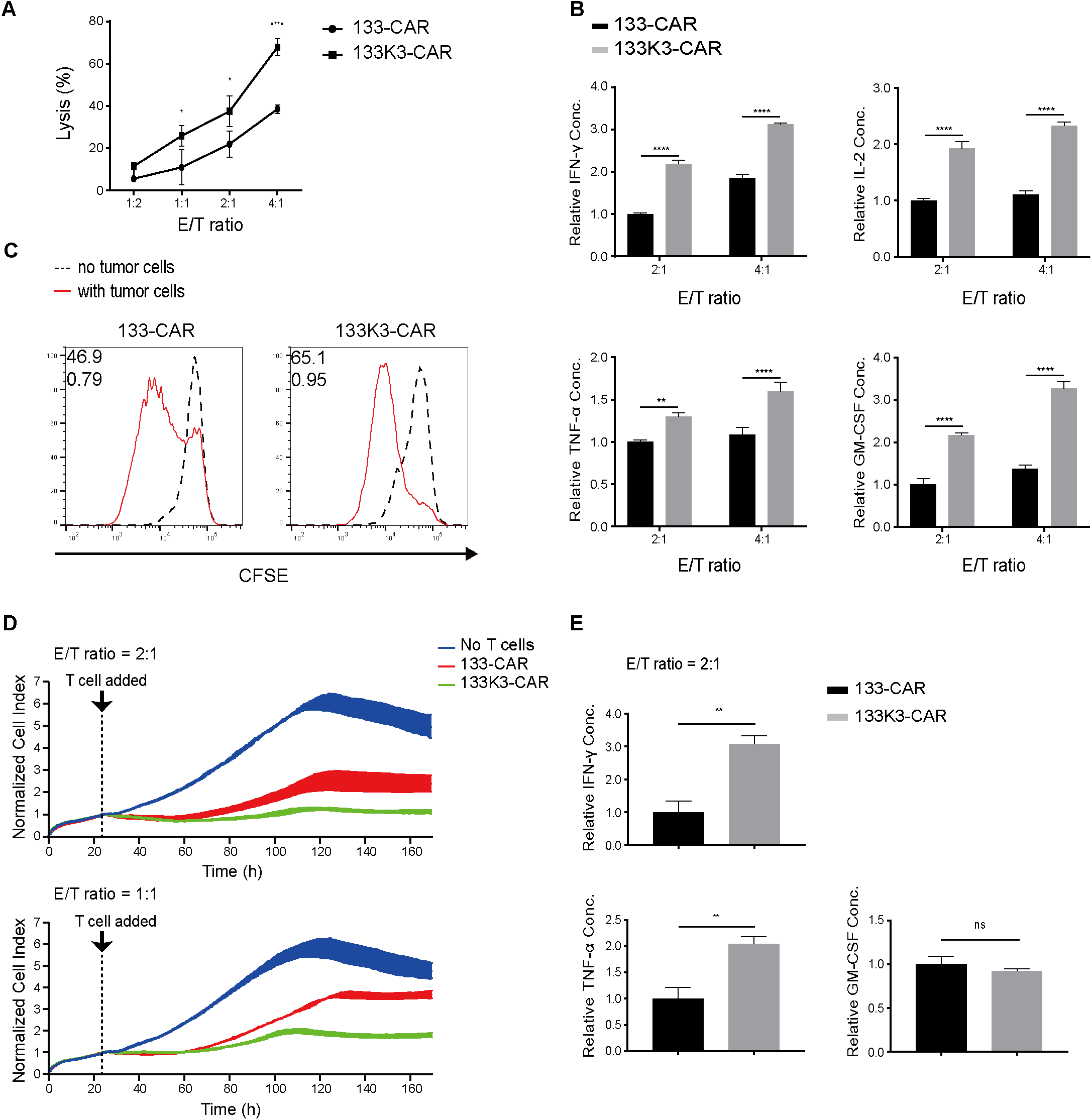
133K3-CAR T cells showed improved antitumor efficacy compared with 133-CAR T cells *in vitro*. (A-E) Same as Fig. 2A-E except that the tumor cells were U251 cells expressing firefly luciferase and CD133, and T cells expressed CD133-specific CAR.

## Conclusion

*IKZF3* KO in CAR T cells can markedly enhance their therapeutic effects in treating different types of solid tumors, suggesting that *IKZF3* KO is an attractive alternative to the use of LENA in enhancing CAR T functions.

## Materials and Methods

### Plasmids

*PiggyBac* transposon vector expressing CD133-specific CAR and the plasmid expressing *piggyBac* transposase (PBase) were prepared according to Zhu et al^45^. HER2-specific CAR expression construct was generated by replacing CD133-specific scFv with the HER2-specific scFv derived from the mAb Herceptin. The Cas9 expression plasmid pST1374-Cas9-GFP was created according to Su et al^38^. The sgRNA pair targeting the *IKZF3* gene locus was designed using E-CRISP website (http://www.e-crisp.org/E-CRISP/) and expressed from pGL3-U6 sgRNA-PGK-Puro vector (Addgene 51133).

### Tumor cell lines

Glioma U251 and breast cancer MDA-MB-453 lines were purchased from Cell Bank of the Chinese Academy of Sciences (Shanghai, China) and cultured in DMEM containing 10% FBS and 1% penicillin/ streptomycin. The cells were routinely checked for mycoplasma infection. To generate the U251-CD133OE line, the *piggyBac* transposon vector (2.5 μg) expressing the human CD133 antigen^45^ was co-delivered with a PBase expression vector (2.5 μg) into 2 ⨯ 10^6^ U251 cells using Nucleofector 2b (Lonza, Köln, Germany), followed by puromycin (500 ng/ml) selection of the stably transfected cells for 14 days. To generate the U251-CD133OE-ffluc and 453-ffluc lines, 2 ⨯ 10^6^ U251-CD133OE and MDA-MB-453 cells were nucleofected with a *piggyBac* transposon vector (2.5 μg) expressing firefly luciferase with the PBase vector (2.5 μg) and the stably transfected cells selected on G418 (500 μg/ml) for 14 days.

### CAR-T cell generation and expansion

PBMCs from healthy donors were purchased from HemaCare (PB009C-3). To generate IKZF3-deficient CAR-T cells, 15-20 million PBMCs in RPMI 1640 (10% FBS, 1% penicillin/streptomycin) were nucleofected using Nucleofector 2b (Lonza, Köln, Germany) with a mixture of 4 plasmids (5 μg each) expressing CAR, PBase, gRNAs and Cas9, respectively. IKZF3-sufficient control CAR-T cells were generated in the same way except that the gRNAs and Cas9 expression vectors were omitted during nucleofection.

24 h following nucleofection, a mixture of allogenic PBMCs derived from 5 donors, irradiated at a dose of 40 Gy with an X-ray irradiator (Rad Source Technologies), were added to the nucleofected cells at the ratio of 15:1 in the presence of anti-CD3 antibody (50 ng/ml, 130-093-387, Miltenyi Biotec) and IL-2 (300 IU/ml, Novoprotein) as described ^45^. 48 h later, the culture media was replaced with AIM V containing 10% fetal bovine serum (Gibco) and 300 IU/ml IL-2 (Novoprotein). Puromycin (0.5 μg/ml) was added and CAR-expressing T cells selected for at least 7 days. CAR expression and activation/differentiation of the expanding T cells were monitored by flow cytometry (CytoFLEX cytometer, Beckman Coulter) twice a week (see further). 14 days after nucleofection, the expanded CAR T cells were re-stimulated with PBMC and anti-CD3 as in the first-round stimulation and further expanded for 10 days.

### Flow cytometry

CAR-T cells were stained with anti-Myc antibody (clone 9B11, 2276S, Cell Signaling Technology) to detect Myc-tagged CAR, with anti-CD45RO (559865, BD Biosciences) and anti-CD62L (562719, BD Biosciences) to evaluate differentiation into memory and effector cells, and with anti-CD69 (555533, BD Biosciences) to assess activation. CD133 and HER2 expression on the tumor cell lines were detected using anti-CD133 (130-098-826, Miltenyi Biotec) and anti-HER2 (BMS120Fl, ThermoFisher Scientific), respectively. All samples were washed with FACS buffer (PBS with 0.5% BSA and 2 mM EDTA) before addition of antibodies. After incubation in 4°C for 30 min, cells were washed with FACS buffer prior to analysis on the flow cytometer. We collected the data on 10000 events and in all cases appropriate isotypes were used as negative controls. Flow cytometry was performed on a CytoFLEX cytometer (Beckman Coulter), and analyzed using FlowJo software.

### DNA extraction, genotyping and sequencing

Genomic DNA was extracted using DNeasy® Blood & Tissue Kit (QIAGEN) before PCR amplification of the targeted site at the *IKZF3* locus using the following primers: Forward GAGAACCCTTCCTCTCCCCT; Reverse ACGTGGCTGCATTAGGAGAG The resulting PCR amplicons (658 bp) were cloned into pGEMT vector (Promega) before Sanger sequencing of individual inserts.

### Western blot analysis of IKZF3 expression

2 ⨯ 10^6^ CAR T cells collected after 24-day expansion were resuspended with 1 ⨯ Laemmli sample buffer (BIORAD) containing 2.5% β-mercaptoethanol before sonication with 50% amplitude in Sonic Dismembrator (FisherBrand). The lysate was heated at 95°C for 5 min and centrifuged at 10000 rpm/s for 2 min to remove the insoluble material prior to SDS-PAGE in a 4-20% gradient ExpressPlus^™^ PAGE Gels (Genscript) at constant 120 V for 1 h. The proteins were transferred to PVDF membranes with Semi-Dry Electrophoretic Transfer Cell (BIORAD) at constant 15V for 30min. The membranes were blocked by 5% BSA in PBS with 0.05% Tween-20, followed by incubation with rabbit anti-IKZF3 (NPB2-24495, Novus) or mouse anti-GAPDH (sc-32233, Santa Cruz) antibodies in 4°C overnight. Membranes were then washed in PBST, followed by 1h incubation with HRP-conjugated donkey-anti-rabbit antibody (711-035-152, Jackson) or donkey-anti-mouse antibody (711-035-150, Jackson) for IKZF3 or GAPDH protein, respectively. After washing in PBST, the proteins were visualized with Chemiluminescent HRP Substrate (Merk Millipore) and signals recorded using Bio-Rad ChemiDoc MP chemiluminescence imaging system (BIORAD).

### Luciferase-based cytolytic assay

Firefly luciferase-expressing tumor cells were plated in triplicates in opaque white walled plate (Corning), and CAR T cells were added according to desired effector to target ratios. 48 h (for HER2-specific CAR T) or 72 h (for CD133-specific CAR T cells) later, D-Luciferin potassium salt (PerkinElmer) dissolved in sterile water was added at the final concentration of 150 μg/ml. The Bioluminescence signal was recorded using the EnVision® Multimode Plate Reader (PerkinElmer) and the percentage of tumor lysis was calculated according to the following formula: Tumor cells lysis (%) = 100% × (1 – signal _co-culture experiment_ / signal _tumor alone_).

### Impedance-based real time cytolytic assay

The cytolytic capability of CAR T cells over a 180-h period was monitored using xCELLigence RTCA system (ACEA Biosciences). MDA-MB-453 or U251-CD133OE were plated in E-plate (ACEA Biosciences) in quadruplicate at 5000 or 10000 cells per well, respectively. After 24 h, CAR T cells were added according to desired effector to target ratios. Lysed tumor cells detached from the bottom of the electrode plate and changed electrical impedance, which was recorded by the system. The impedance-based cell index was measured in 15 min intervals, and individual cell index was normalized with the cell index prior to the addition of CAR T cells.

### Cytokine secretion assay

Tumor cells and CAR T cells were co-cultured according to desired effector to target ratios in triplicates for 24 h. The supernatant was collected and various cytokines measured using the PerkinElme AlphaLISA kit (IL-2, AL221C; IFN-γ, AL217C; TNF-α, AL208C; GM-CSF, AL216C).

### T cell proliferation assay

1-3×10^6^ HER2-specific CAR T cells were labeled with 0.5 μM CFSE (eBioscience) and cultured with equal numbers of irradiated MDA-MB-453 cells (70 Gy via X-ray irradiator, Rad Source Technologies) for 7 days in RPMI containing 10% FBS, 1% penicillin/streptomycin. 1 ⨯ 10^5^ cells were analyzed for CFSE signal by flow cytometry before and at the end of culture. The data were analyzed by Flowjo to determine the Percent Divided, the value for the percentage of cells that divided at least once, and the Division Index, the average number of divisions for all cells in the original population. The proliferation of CD133-specific CAR T cells was analyzed in the same way except that half the amounts of U251-CD133OE cells were used and the cells cultured for 4 days.

### Xenograft model and tumor growth *in vivo*

Animal experiments were performed at the National Center for Protein Science in Shanghai following the guidelines of the Shanghai Administrative Committee for Laboratory Animals. 1.5 ⨯ 10^6^ 453-ffluc cells in 100 μl sterile PBS were mixed with equal volume of ice-cold Matrigel® (Corning). 200 μl of the mixture was subcutaneously injected over the dorsal flank of the right foreleg into 6-week-old NPSG females (NOD-Prkdc^scid^ Il2rg^null^, Shanghai Jihui Laboratory Animal Care Co.,Ltd.). Six days later, tumor-bearing mice were randomly grouped and intravenously injected with HER2-CAR, HER2K3-CAR T cells (each at 5 ⨯ 10^6^ cells in PBS per mouse) or PBS. Tumor growth was monitored by measuring the tumor volumes and imaging. Tumor volumes, defined as (length × width^2^)/2, were determined using a caliper. To measure the bioluminescence signal, the mice were anesthetized via administration of 2% isoflurane with Matrx VIP 3000 Calibrated Vapoizer (MIDMARK) at 2 L/min flow rate of total input oxygen. D-Luciferin potassium salt (PerkinElmer) was injected intraperitoneally (150 mg/kg), and tumors imaged with the IVIS Lumina II device (PerkinElmer) 10 min afterwards. The imaging was performed twice a week 6 days after the subcutaneous inoculation of tumor cells. The mice were sacrificed when body weight loss reached 20% or the tumor volume reached 1000 mm^3^.

### RNA-seq

After expansion and selection for 24 days, 1 ⨯ 10^6^ *IKZF3* knockout and parental HER2-specific CAR T cells were collected. Total RNA was isolated by phenol-chloroform extraction method and its integrity evaluated using the Agilent 2100 Bioanalyzer (Agilent Technologies). The libraries were constructed using TruSeq Stranded mRNA LTSample Prep Kit (Illumina), followed by sequencing on the illumine sequencing platform (HiSeqTM 2500, Illumina, Shanghai OE Biotech Co., Ltd.), and 150-bp paired-end reads were generated. All the raw data were checked with fastQC (https://www.bioinformatics.babraham.ac.uk/projects/fastqc/) for quality control. Fastq files were processed with Salmon^46^ and the reads were mapped to human hg38 transcriptome. The data was subsequently uploaded into iDEP (http://bioinformatics.sdstate.edu/idep/) to analyze differentially expressed genes (DEGs). Enrichment analysis for the DEGs was performed using DAVID Bioinformatics Resources 6.8 (https://david.ncifcrf.gov/). The RNA sequencing data has been deposited in the NCBI database with accession number PRJNA684699.

## Acknowledgements

The authors would like to thank the HTS platform in SIAIS and the Molecular and Cell Biology Core Facility (MCBCF) at the School of Life Science and Technology, ShanghaiTech University, and the Animal Facility at the National Facility for Protein Science in Shanghai (NFPS), Zhangjiang Lab, Shanghai Advanced Research Institute, Chinese Academy of Science for providing technical support. This work was supported by the National Key R&D Program (2019YFA0111001) of China.

## Supplemental Information

Table 1 contains sequence of primers for PCR amplification of targeting sites.

An excel file listing all the differentially expressed genes (FC>=1.5, q<=0.1) is included.

## Author contributions

Y.Z., T.C. and X.Z. designed the research. Y.Z. performed the experiments. Y.Z., T.C. and X.Z. analyzed the data. Y.Z., T.C., Z.J. and Q.Y. performed the RNA sequencing analyses. L.L. and J.T. coordinated the research. Y.Z. and T.C. wrote the manuscript. T.C., X.H. and X.Z. revised the data and performed a final revision of the paper.

---

**Fig. S1.**
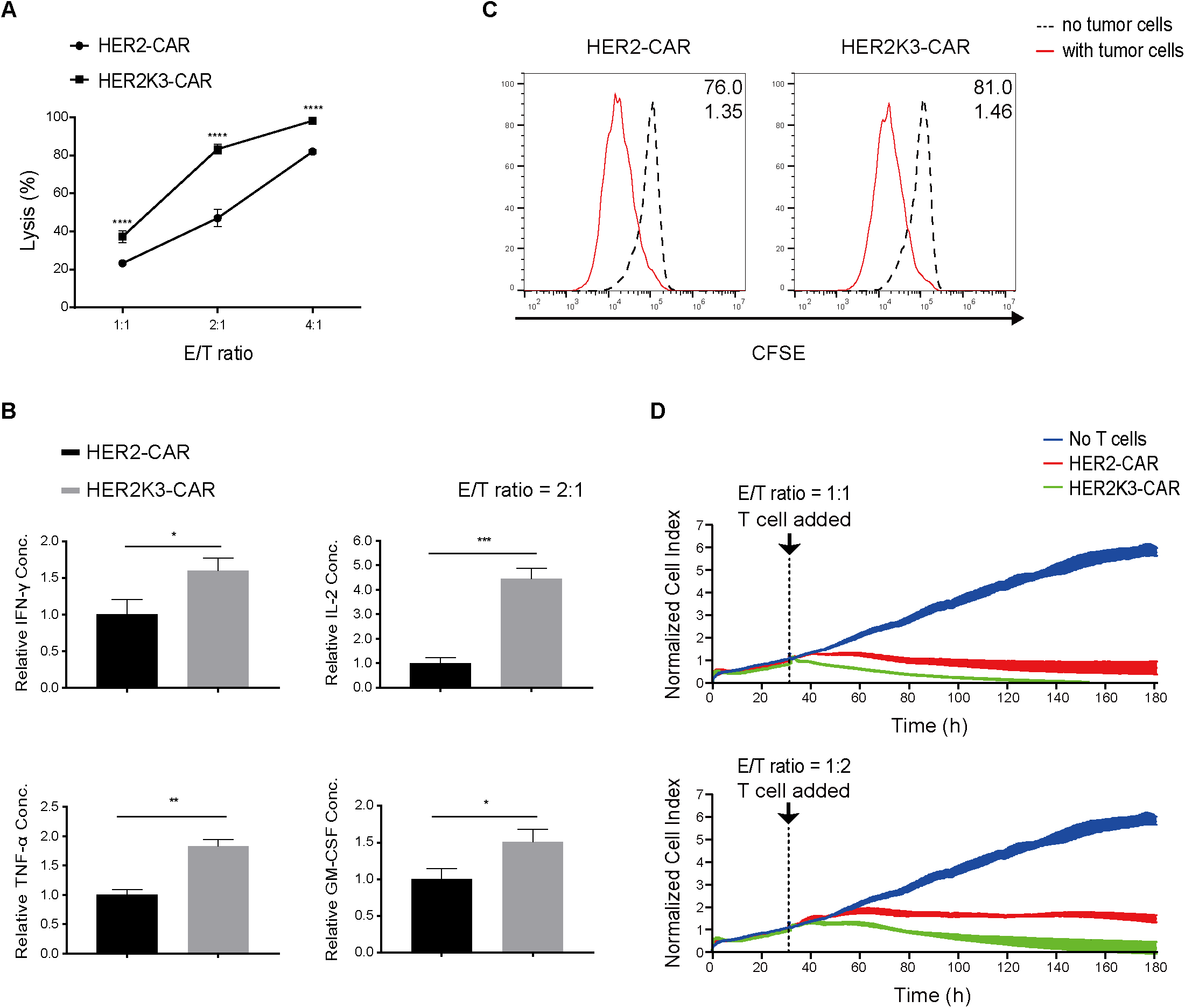
*IKZF3* KO enhances HER2-CAR T activity *in vitro*. Shown is a biological replicate for the experiment described in Fig. 2A-D, except that in Fig. S1B, only one E/T ratio was tested.

**Fig. S2.**
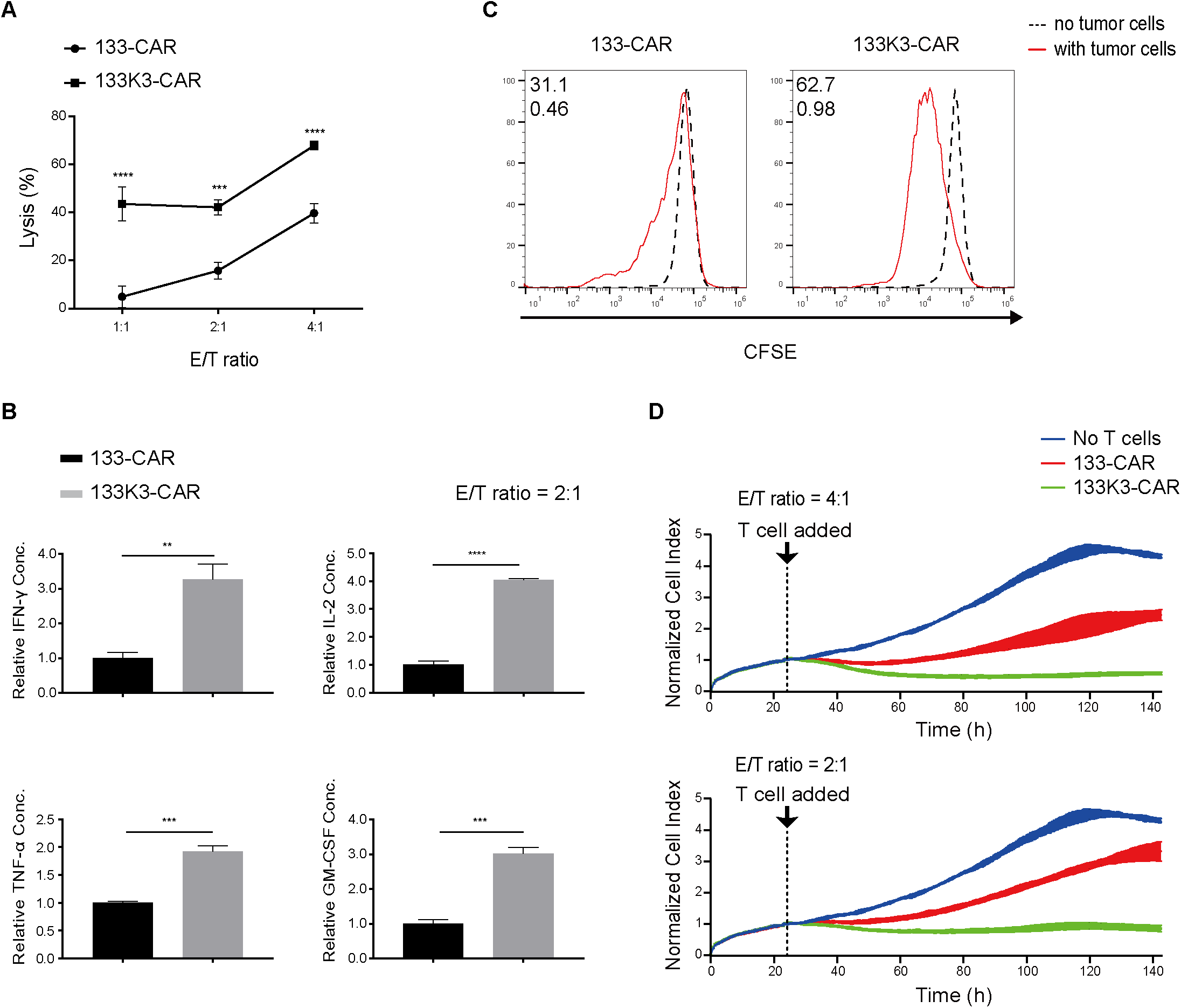
F133K3-CAR T cells showed improved antitumor efficacy compared with 133-CAR T cells *in vitro*. Shown is a biological replicate for the experiment described in Fig. 6A-D except that in Fig. S2B, only a single E/T ratio was tested.

